# Mutation breeding for quantitative traits in the era of genomic selection

**DOI:** 10.64898/2026.10.27.762151

**Authors:** Chin Jian Yang, Rajiv Sharma, Ian Mackay, Wayne Powell

## Abstract

Mutation breeding creates new genetic variation by inducing random mutations and screening for mutants with desirable phenotypes, whereas selective breeding exploits variants that already segregate in breeding populations. The two approaches have rarely been combined, and whether genomic selection (GS) can capture polygenic variation induced by mutagenesis has not been tested. We evaluated GS in two public panels of advanced mutants (190 wheat and 74 tomato lines) and in simulated biparental and multiparental populations derived from mutants. In both panels, predictive abilities (up to 0.39 in wheat and 0.45 in tomato) came entirely from variants shared among lines, which in wheat formed chromosome segments segregating from the parental cultivar. Variants private to single lines, where induced mutations lie, carried no predictive signal. In simulated crosses in which mutations segregate, GS outperformed marker-assisted selection (MAS) of the largest mutation when mutations were numerous and their effects balanced (genetic value 2.84 vs 0.45 after two cycles), but not when unfavorable mutations dominated. In simulated multiparental populations, mutant founders increased genetic gain over five cycles of GS by 1.3- to 6.8-fold when mutational variance was large, but by only 4–5% when it was small relative to standing variation. Induced polygenic variation therefore becomes accessible to GS once mutations are shared among relatives through crossing, which offers a route to integrate mutation breeding into pre-breeding programmes.

**KEY MESSAGE:** Genomic selection cannot predict private induced mutations across independent mutant lines, but captures polygenic induced variation once mutants are crossed, provided mutational variance is substantial.

## INTRODUCTION

Plant breeding plays an important role in agricultural innovation by directly contributing to the development of crop varieties with enhanced yield, disease resistance, and adaptability to diverse environments (Borlaug 1983, Bradshaw 2017). Since the early days of domestication, breeding practices have evolved from incidental selection by farmers to more systematic approaches involving statistics, experimental design, biotechnology, multi-omics and precision breeding technologies (Wallace et al 2018, Bernardo 2020). Artificial selection in modern plant breeding primarily targets naturally occurring genetic variants and haplotypes that arise from spontaneous mutations and survive through natural selection, genetic drift, and recombination over generations (Huang and Han 2014). Different breeding strategies, such as line, hybrid, and clonal breeding, are tailored to the biological characteristics of a given species including self-compatibility, hybrid vigor, generation time, and ploidy levels (Acquaah 2015). Key applications in plant breeding include genomic selection (GS) (Desta and Ortiz 2014), marker-assisted selection (MAS) (Ribaut and Hoisington 1998), phenomic selection (Rincent et al 2018), gene editing (Chen et al 2019), doubled haploid (DH) (Gilles et al 2017) and tissue culture (Thorpe 2007). Among these, GS stands out for its ability to improve selection for quantitative traits through accurate prediction of breeding values. As the field progresses, plant breeding will continue to play a critical role in enhancing crop resilience to climate change and ensuring global food security (Xiong et al 2022, Cooper and Messina 2023).

Mutation breeding is a specialized form of plant breeding that induces novel genetic variants using chemical or physical mutagens. However, the history of mutation breeding traces back to selection for spontaneous mutations in rice breeding in the early 20^th^ century when formal mutagenesis protocols were not yet available (Suganami et al 2024). Nowadays, instead of relying on spontaneous mutations, which occur at extremely low heritable mutation rates of 10^-11^ to 10^-8^ per base pair (Ossowski et al 2010, Quiroz et al 2023), mutation breeding raises mutation densities by several orders of magnitude, to around 10^-6^ per base pair using mutagens (about 1.5 to 4.5 mutations per Mb in EMS-treated barley; Huang et al 2023). This approach is particularly beneficial for crops with limited genetic diversity or those that are not quite amenable to conventional breeding strategies. While the specifics of mutation breeding programs can vary across crops and breeders, the general process involves mutagenesis and dosage optimization, followed by phenotypic screen and variety development, sometimes with mutation mapping (Figure 1). Common mutagens used include ethyl methanesulfonate (EMS) (Chen et al 2022), gamma rays (Ahumada-Flores et al 2021, Jankowicz-Cieslak et al 2022), X-rays (Jankowicz-Cieslak et al 2022), ion beams (Kazama et al 2011, Hase et al 2012) and cosmic rays combined with low gravity in space (Mohanta et al 2021). Optimizing the mutagen dosage is critical to generate sufficient number of viable mutants with novel phenotype (Muhammad et al 2021, Chakraborty et al 2023). Mutagenesis is typically conducted on wild type seeds, though it can also be performed on whole plants and pollen (Oladosu et al 2015, FAO/IAEA 2018, Settles 2020). Because large effect mutations are often recessive, phenotypic screen generally occurs in the M2 generation, but exceptions exist for screening in M1 generation when chromosome aberrations such as large insertions, deletions or aneuploidy are involved. Once a line exhibiting a desirable mutant phenotype is identified, it can be crossed to its wild type to eliminate background mutations or to map causal mutations using bulked segregant analysis (Abe et al 2012, Fekih et al 2013). In many cases, a classical Mendelian segregation ratio in the F2 indicates that a single gene is responsible for the observed mutant phenotype. Mutation breeding has been successfully applied to a variety of crop species, including field crops (Lethin et al 2020), ornamental plants (Yamaguchi 2018, Melsen et al 2021) and fruit trees (McCord et al 2022). Overall, mutation breeding is a valuable tool in plant breeding for creating novel genetic diversity and developing improved crop varieties (Sivasankar et al 2021).

**Figure 1.**
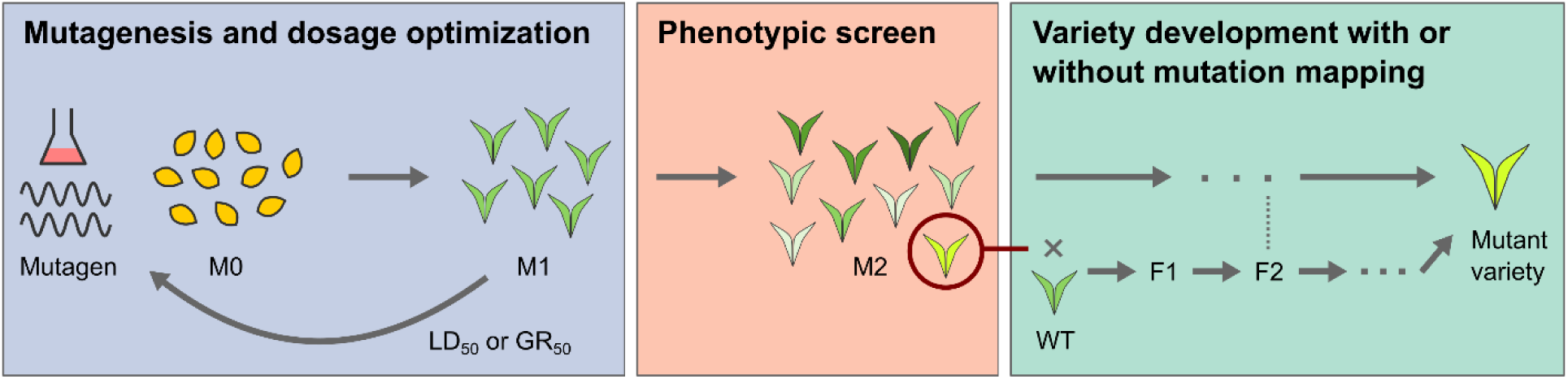
General pipeline in mutation breeding programs. Abbreviations: Mutant generation 0/1/2 (M0/M1/M2), median lethal dose (LD50), median growth reduction (GR50), wild type (WT).

Outside of mutation breeding, spontaneous and induced mutations are extensively studied in population and evolutionary genetics (Loewe and Hill 2010). While a comprehensive overview of the field is beyond our scope, we highlight relevant aspects that may be of interest to mutation breeders. Polygenic mutations, which are mutations of small effects (Lynch 1988), can be pervasive (Mukai 1979) but have historically received less attention than large effect mutations. Polygenic mutations contribute to phenotypic variation in traits in a resembling role of polygenic natural variants in quantitative traits. Predicting the distribution of additive genetic effects in polygenic mutations is complex, as it varies with mutation sources, species and traits. Oka et al (1958) and Mukai et al (1984) showed in *Oryza sativa* (rice) and *Drosophila melanogaster* (fruit fly) that polygenic mutations result in populations with a phenotypic mean of zero, indicating non-directionality with equal likelihood of beneficial and deleterious mutations. Similarly, Shaw et al (2002) observed equal proportions of beneficial and deleterious mutations in *Arabidopsis thaliana* (thale cress), but Keightley and Lynch (2003) proposed that most mutations are deleterious. In contrast, Bondel et al (2019) found an overall excess of beneficial mutations but a tendency for large effect mutations to be more deleterious than beneficial in *Chlamydomonas reinhardtii* (green alga). Keightley and Ohnishi (1998) further showed that most mutations in *D. melanogaster* have deleterious effects across multiple traits. A review by Eyre-Walker and Keightley (2007) suggested that deleterious mutations are generally more common across species than beneficial ones. Regardless of the directionality of mutation effects, these effects can be modeled using a reflected gamma distribution, where positive and negative effects are represented by two separate gamma distributions with the proportion of positive effects specified (Keightley and Ohnishi 1998, Keightley and Eyre-Walker 2010). In terms of dominance, the effects are thought to resemble a metabolic model, where greater degrees of dominance are associated with larger additive effects while smaller additive effects exhibit lower dominance (Phadnis and Fry 2005). Despite this wealth of knowledge, there remains a disconnect between population geneticists and mutation breeders. As Sharma et al (2025) suggests, it is timely to bridge these fields and leverage interdisciplinary expertise to address real world challenges.

Here, we seek to bridge conventional selective breeding and mutation breeding by developing a unifying approach that combines the best techniques from both disciplines. We evaluate the feasibility of applying GS in estimating mutation effects and breeding values for two different purposes. First, GS can be used to screen higher number of mutants, improving the odds for identifying useful mutants. Second, GS can be used to estimate mutation-specific breeding values for germplasm enhancement in breeding programs using novel mutations. Our workflow is briefly described here. We choose two publicly available datasets of mutant populations in wheat (Guo et al 2019) and tomato (Shikata et al 2016, Yano et al 2019), and simulate biparental and multiparental populations involving mutants as our test cases for each area of application. For the two mutant populations, in addition to using a Single Nucleotide Polymorphism (SNP) based genomic markers, we also collapsed the SNP markers according to genes and Gene Ontology (GO) annotations. Critical assumptions on the distribution of mutation rate, number and effect in our simulation are based on the current literature in mutation breeding and population genetics (Keightley and Ohnishi 1998, Phadnis and Fry 2005, Keightley and Eyre-Walker 2010, Yano et al 2019, Huang et al 2023). Even though dominance is an important consideration in mutation studies, dominance effects are not considered here because selection candidates are inbred (doubled haploid) lines, in which only additive effects are transmitted and expressed. For each population and trait, we apply the GBLUP model (VanRaden 2008) and compute the prediction accuracy using a standard five-fold cross validation approach (Werner et al 2020). For the two simulated populations, we also compare the genetic values and mutation allele frequencies under recurrent selection with GS against conventional strategy of MAS with the largest favorable mutation.

Our results show where GS can and cannot help mutation breeding. Across independent lines of an advanced mutant panel, prediction relies on variants shared among lines, which in the wheat panel were background chromosome segments rather than induced mutations, while private induced variants carry no predictive signal. Once mutations segregate in crosses, GS captures polygenic induced variation: in simulated biparental populations it outperformed MAS when mutations were numerous and of mixed effect, and in simulated multiparental populations mutant founders increased genetic gain when mutational variance was large relative to natural variance. The concept was recently outlined in a review on mutation breeding and genomic selection (Bhatnagar-Mathur et al. IAEA, personal communication); here we test it empirically and by simulation.

## MATERIALS AND METHODS

### Advanced mutant populations

The dataset for a wheat mutant population was obtained from Guo et al (2019), which contained 190 mutants at advanced (post-M2) stage and 10 traits evaluated in 5 environments. Of these 190 mutants, 181 and 9 were developed from EMS and gamma-ray mutagens, respectively. All of the mutants have been genotyped using the wheat 660k array (Sun et al 2020) and 67,402 high quality SNP markers were publicly available (Guo et al 2019). The 10 traits included plant height (PH), spike length (SL), thousand grain weight (TGW), maximum tiller numbers (MT), effective tiller numbers (ET), pre-winter tiller numbers (PWT), grain number per spike (GNS), flag leaf angle (FLA), spikelet number per spike (NSL) and spikelet density (SD). For each trait, best linear unbiased estimates (BLUEs) were calculated for each mutant line using fixed line effect and random environment effect in R/sommer package (Covarrubias-Pazaran 2016). This dataset was previously used in Genome-Wide Association Studies (GWAS) and 32 large effect mutations were mapped (Guo et al 2019).

The dataset for a tomato mutant population was obtained from Shikata et al (2016) and Yano et al (2019), which contained 74 mutants at advanced (M3 to M7) stage and 20 traits were evaluated in one environment. Of these 74 mutants, 70 and 4 were developed from EMS and gamma-ray mutagens, respectively. All of the mutants have been genotyped using whole genome sequencing and 175,451 high quality genomic markers were publicly available (Yano et al 2019). The 20 traits included fruit weight, Brix value, carotenoids, and levels of amino acids (Asp, Thr, Ser, Asn, Glu, Gln, Gly, Ala, Ile, Leu, Tyr, bAla, Phe, GABA, His, Lys and Arg). For each trait, BLUEs were calculated for each mutant line as simple means of the plants with data (up to three plants per line; on average 2.8 plants with non-missing values). This dataset was previously used in genomic characterization of mutational landscape and a single large effect mutation responsible for curly leaf phenotype was mapped (Yano et al 2019).

### Variant burden and quality control

For each mutant population, the variant burden of each line was calculated relative to the wild type (WT). In wheat, the WT was represented by two genotyped WT samples; only SNPs with concordant calls in both samples (67,309 of 67,402) were used, and the burden of a line was the proportion of these SNPs at which its genotype differed from the WT. Lines differing from the WT at more than 5% of SNPs or with more than 5% heterozygous calls were classed as off-types (outcrosses or seed admixture) and removed from the quality-controlled (QC) set. In tomato, the genotype data consist of variant calls, so the burden was the number of variant sites carried by each line. In both species, each non-WT site was classed as private (carried by one line) or shared (carried by two or more lines). Independent induced mutations are expected to be almost entirely private, so shared sites indicate pre-existing variation. In wheat, linkage disequilibrium (r) between physically adjacent shared SNPs was compared with that between random pairs of shared SNPs to test whether shared SNPs form chromosome segments.

### Population structure

For each population, the population structure was characterized using a Principal Component Analysis (PCA). If mutations are truly random, the population structure should be absent. PCA was calculated by first computing the additive genetic relationship matrix (K) using R/sommer. Principal components were then calculated using the eigen() function in R. The first two eigenvectors or principal components were plotted along with their percent variance explained as given by the eigenvalues.

### GS in advanced mutant populations

Within each population and trait, the following mixed model was fitted using R/sommer (Covarrubias-Pazaran 2016) to obtain the genomic Best Linear Unbiased Predictions (gBLUP) for random genetic effect, which is also known as genomic Estimated Breeding Values (gEBV). Predictive abilities were computed using five-fold cross-validation repeated 20 times with different random partitions (Gianola and Schön 2016). This procedure involved random partitioning of mutant lines into five approximately equal groups, using four groups to train the GS model for deriving gEBV in the fifth group, repeating the model fitting step for all possible group combinations, and estimating prediction accuracies as the correlations between gEBV and observed phenotypic trait values.

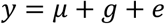

*y* is a vector of phenotypic trait values for mutant lines,

*μ* is a vector of fixed intercept effect,

*g* is a vector of random genetic effect with an assumed distribution of *N*(0, **K**σ^2^_g_),

*e* is a vector of random residual effect with an assumed distribution of *N*(0, **I**σ^2^_e_).

Predictive ability is reported as the mean and standard deviation across the 20 repeats. Folds in which the REML estimate of the genetic variance was zero were recorded as missing. In addition to all markers, genomic relationship matrices were built from (i) shared and (ii) private non-WT sites, and from markers collapsed into genes and GO categories (biological process, cellular component and molecular function), treating any variant in a gene, or in any gene annotated to a GO term, as one allele. As a structure-only benchmark in wheat, test lines were predicted by the training-set mean of their group (off-type or true-to-type). For the repeated cross-validation, GBLUP was fitted by REML with R/rrBLUP (Endelman 2011), which gave the same predictive abilities as R/sommer on these data.

### Simulated biparental population

All populations were simulated based on tomato, which is an important diploid crop species with 12 chromosomes. Genetic map lengths from a tomato multiparental population (Pascual et al 2015) were used as the foundation for simulation. The lengths ranged from 121 to 215 centiMorgans (cM) with a total of 2,154 cM. Crosses and recombination events were simulated using R/AlphaSimR (Gaynor et al 2021). Trait genetic architectures including effects and numbers of genomic variants due to natural and mutation sources were simulated using custom R scripts. GS on simulated individuals were done using gEBV obtained from fitting Ridge Regression BLUP (RRBLUP) mixed models in R/rrBLUP (Endelman 2011). While both RRBLUP and gBLUP models are equivalent (VanRaden 2008), RRBLUP was a more convenient option in this case as the RRBLUP marker effects could be directly used to calculate gEBV in subsequent generations without fitting the models repeatedly.

The biparental population was initiated by crossing two inbred parents, a Wild Type (WT) and an advanced mutant line, followed by a Doubled Haploid (DH) induction to create 200 inbred lines. The advanced mutant line originated from the same WT, so they share the same genetic background except for induced mutation sites. For convenience, the genotyping method was assumed to capture only one mutation in every 0.1 cM, which would equate to a maximum of 21,540 possible genotyped mutations. Given this assumption, the numbers of mutations or genetic markers were set to either 10 or 1,000 to represent low (5e^-4^) and high (5e^-2^) mutation rates. These mutations were sampled uniformly across the whole genome. Every mutation was assumed to be genotyped and no non-causal markers were included, which represents a best case for GS. Proportions of favorable mutations were set to either 0.1 or 0.5 to represent low and equal numbers of mutations with desired direction of effects. Additive genetic effects were simulated based on a gamma distribution with shape and scale parameters of 0.5 and 1 respectively (Keightley and Ohnishi 1998). Unfavorable effects were sampled uniformly according to the proportions by setting those effects to negative. Because advanced mutant lines would have undergone some selection pressure during mutant screen, the largest effect was reserved as favorable. The additive genetic effects were scaled to produce a genetic variance of 1. Genetic values of the simulated DH lines were obtained by totaling their mutation effects and phenotypic trait values were calculated as the sums of genetic values and residual values drawn from a random normal distribution with a variance of 1. Therefore, the heritability of the simulated trait was 0.5.

The performance of GS method with two cycles of recurrent selection was compared against the baseline using a conventional MAS method. In cycle 0, the RRBLUP model was trained using an initial set of 200 DH lines (cycle 0), 10 lines with the best phenotype were selected, 20 random crosses were made among the selected lines, and 10 new DH lines were created from each cross which resulted in a new population with 200 DH lines. In cycle 1, 10 lines with the best gEBV estimated using the previously trained RRBLUP model were selected and the same crossing approach was applied. While it is not necessary to create DH in practice, it allowed us to ignore dominance effects in training the GS model. Because of the additional DH step, the GS method required two generations per selection cycle. Genetic values of all DH lines and population frequencies of mutant alleles in each cycle were recorded. Conventional MAS method involved mapping linked or causative markers, selecting several lines carrying the marker with the largest effect, and removing background mutations over multiple generations. For convenience, four generations were assumed sufficient for the background selection, which was completed at the same time as cycle 2 in the GS method. Genetic values for the final lines selected using MAS were recorded, and the largest effect allele was fixed at a frequency of 1 and the rest were fixed at a frequency of 0. The simulation was repeated for 100 iterations. Two reference values were also recorded: the WT (genetic value 0) and an ideal line carrying all favorable mutations and none of the unfavorable ones (twice the sum of the positive effects).

### Simulated multiparental population

Similarly, the multiparental population was simulated using the same parameters from tomato. The multiparental population was initiated by making all 28 possible crosses among the 8 inbred founders, followed by a DH induction to create 560 inbred lines. There were two main types of population simulated here. The first type (Pop 1) used inbred founders with no induced mutations and this scenario served as the baseline for comparison. The second type (Pop 2) used advanced mutant lines derived from the same inbred founders, and therefore the resulting simulated population would segregate for natural and mutation genetic variants. Unlike the biparental population simulation, the genotyping method here only captures the segregating natural genetic variants and not the mutations. This is meant to be an advantage of using linkage disequilibrium between natural and mutation variants in the multiparental population to capture the mutation effects without needing to identify mutations from deep sequencing. Only high mutation rate (5e^-2^) was considered here, and the mutations were randomly sampled uniformly across the whole genome. Proportions of favorable mutations were set to either 0.1 or 0.5 to represent low and equal numbers of mutations with desired direction of effects. Additive genetic variances for natural and mutation effects were set to either 0.1 or 1, giving a total of four permutations of possible natural and mutation genetic variances. Same gamma distribution for additive effects and residual variance were used here. The selection approach for the simulated multiparental population was similar to the simulated biparental population, with a few exceptions described here. The number of crosses was set to 45 instead of 20 to account for all possible pairwise combinations of the 10 selected DH lines in each cycle. Instead of stopping at cycle 2, the recurrent selection continued to cycle 5.

To generate the advanced mutant founders of Pop 2, 1,000 M2-derived doubled haploids were simulated for each founder. The number of mutations per doubled haploid was drawn from a negative binomial distribution with mean 75 (half of the 150 expected in an M2 plant, because doubled haploid induction samples one gamete), mutations were placed at randomly chosen marker positions, and their additive effects were drawn from the reflected gamma distribution described above. Phenotypes were simulated with h² = 0.1, and the best doubled haploid of each founder was retained as the mutant founder. Natural QTL were placed at 1,000 of the 1,345 marker positions, with the founders carrying each QTL allele sampled at random. Natural and mutation effects were scaled to the target variances (V_nat, V_mut) in the initial doubled haploid population. From cycle 2 onward, the GBLUP model was retrained in each cycle using the phenotypes and genotypes of the previous cycle only. In the simulations, GBLUP was fitted by REML with a spectral (EMMA) implementation (Kang et al 2008), which gave estimates identical to R/rrBLUP and R/sommer. Each of the eight scenarios was replicated 100 times.

## RESULTS

### Advanced wheat mutant population

Relative to the WT, the median wheat line differed at 2.5% of the 67,309 SNPs with concordant WT calls, and the distribution was bimodal (Figure 2A). Thirty-four lines differed from the WT at more than 5% of SNPs (29 at more than 20%, up to 73%), and some were up to 43% heterozygous. These levels are far above what chemical or physical mutagenesis can produce at array SNP positions, so we classed these lines as off-types from outcrossing or seed admixture. The off-types account for the strong population structure (the first principal component explains 55.1% of the genomic variance; Figure 2B). Among the 156 true-to-type lines, each line carried a median of 1,394 non-WT SNP calls, of which a median of 27 were private: 97% of non-WT calls were shared with at least one other line, and 3,280 SNPs were carried by 10% or more of the lines. Physically adjacent shared SNPs were in almost complete linkage disequilibrium (median r = 0.99, against 0.19 for random pairs of shared SNPs), so the shared SNPs form chromosome segments segregating from the parental cultivar rather than independent induced mutations.

**Figure 2.**
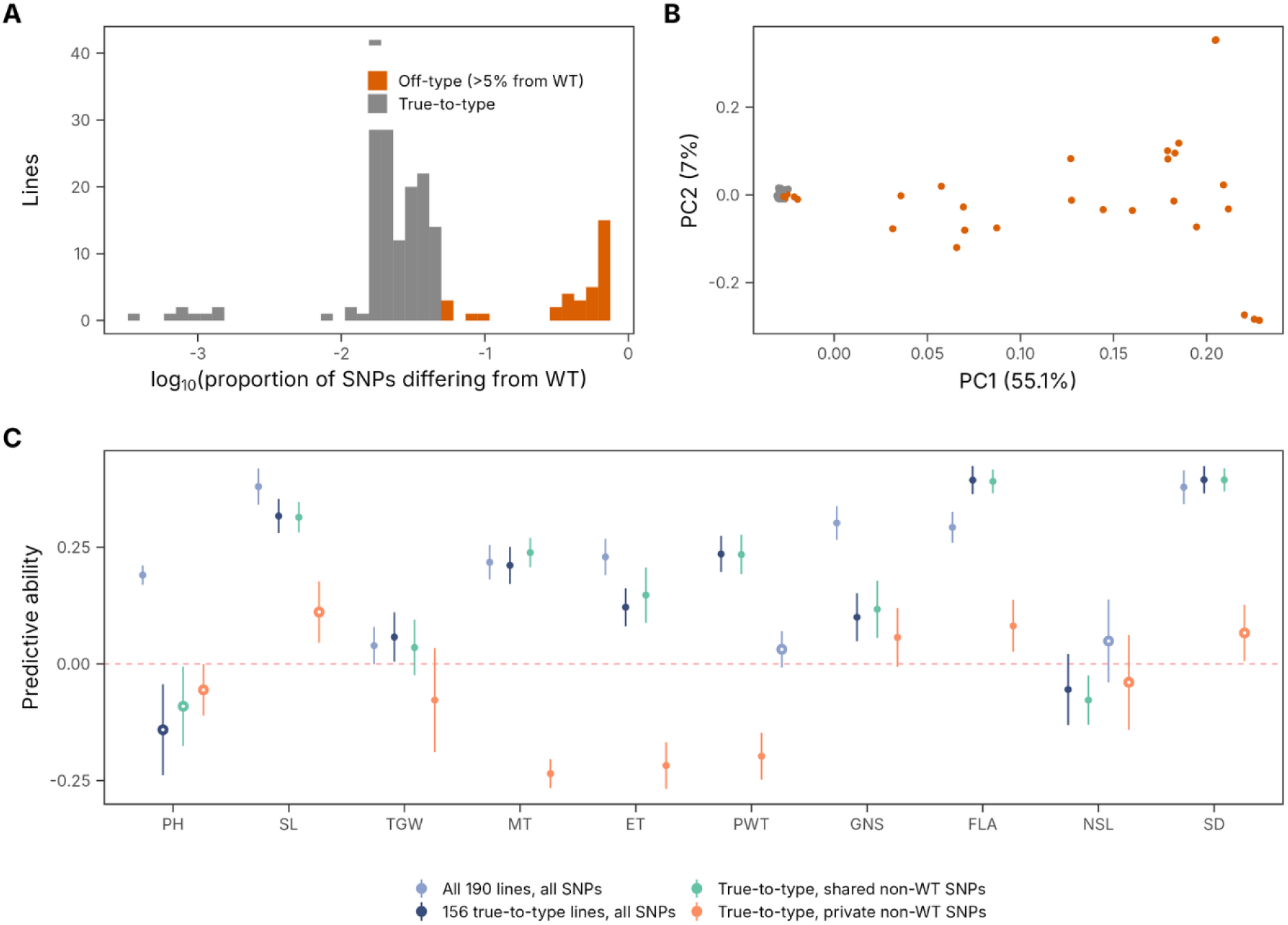
Genomic selection in an advanced wheat mutant population. [A] Proportion of SNPs at which each of the 190 lines differs from the wild type (WT), on a log_10_ scale. Lines differing at more than 5% of SNPs (or with more than 5% heterozygous calls) are classed as off-types. [B] First two principal components of the genomic relationship matrix, coloured by line status. [C] Predictive ability (mean ± SD over 20 repeats of five-fold cross-validation) for all 190 lines with all SNPs, the 156 true-to-type lines with all SNPs, and the true-to-type lines with a relationship matrix built from shared or private non-WT SNPs only. Open circles mark estimates for which the genetic variance was estimated at zero in more than half of the 100 folds.

Using all lines and all SNPs, predictive abilities from 20 repeats of five-fold cross-validation ranged from 0.03 (PWT) to 0.38 (SD) (Figure 2C). Removing the off-types changed the picture trait by trait. Predictive ability for plant height fell from 0.19 to −0.14, whereas FLA (0.39), SD (0.39), SL (0.32), PWT (0.24) and MT (0.21) remained predictable within the 156 true-to-type lines. A model that used only the off-type/true-to-type grouping had no predictive ability (mean r between −0.14 and 0.09 across traits), so prediction was not simply a population-structure effect. However, prediction within the true-to-type lines relied on shared variants. A relationship matrix built from shared non-WT SNPs alone reproduced the full-SNP results (FLA 0.39, SD 0.39, SL 0.31, MT 0.24, PWT 0.23). Private non-WT SNPs had no predictive value: for SL and SD the genetic variance was estimated at zero in 97 and 92 of 100 folds. Collapsing SNPs into genes or GO terms gave no consistent improvement (Figure S1). It raised predictive ability by more than 0.05 only for ET, PH and PWT, and the collapsed markers are dominated by the same shared variants.

### Advanced tomato mutant population

The 74 tomato lines carried a median of 2,412 variant sites each (range 1,453 to 4,226; Figure 3A). Of the 175,451 variant sites, 170,498 (97%) were private to one line and 4,953 were shared. However, 221 sites were carried by more than 10% of the lines, which independent EMS or gamma-ray events would not produce. Population structure was weak: the first principal component explained 2.3% of the genomic variance and mainly separated single lines from the rest (Figure 3B).

**Figure 3.**
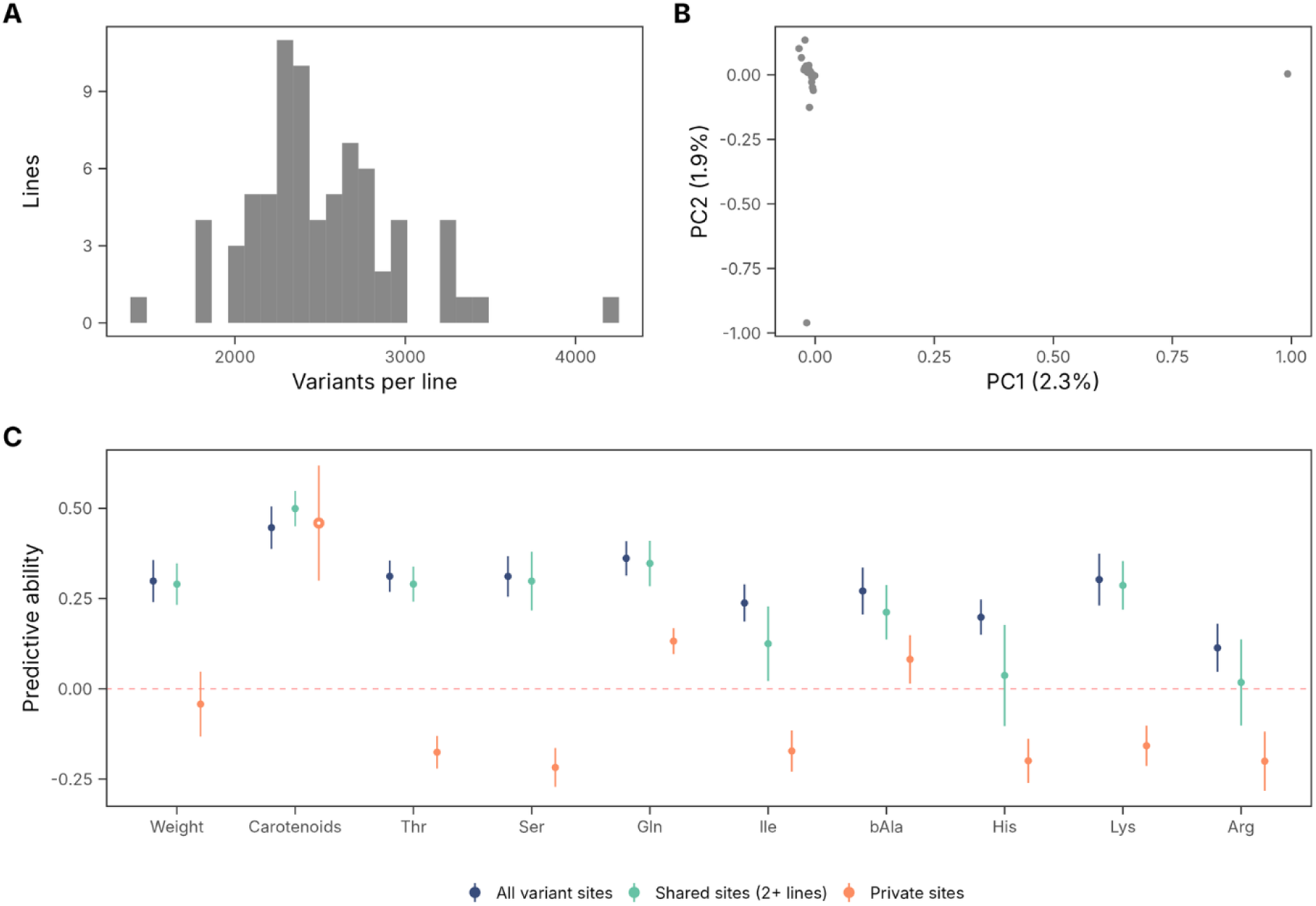
Genomic selection in an advanced tomato mutant population. [A] Number of variant sites per line in the 74 advanced tomato mutants. [B] First two principal components of the genomic relationship matrix. [C] Predictive ability (mean ± SD over 20 repeats of five-fold cross-validation) for ten traits using all variant sites, shared sites (carried by two or more lines) or private sites only. All 20 traits are shown in Figure S2. Open circles mark estimates for which the genetic variance was estimated at zero in more than half of the 100 folds.

Predictive abilities were low to moderate and varied between traits (Figure 3C, S2). With all variant sites, the best-predicted traits were carotenoids (0.45), Gln (0.36), Thr (0.31), Ser (0.31), Lys (0.30) and fruit weight (0.30). For Brix the genetic variance was estimated at zero in all 100 folds. As in wheat, the predictive signal came from shared sites. The 4,953 shared sites alone gave similar predictive abilities (carotenoids 0.50, Gln 0.35, Ser 0.30, weight 0.29, Lys 0.29), whereas the private sites gave values close to zero or negative (Gln 0.13, weight −0.04, Lys −0.16), or a genetic variance of zero in almost all folds (carotenoids, 98 of 100). Collapsing variants into genes or GO terms did not improve prediction (Figure S2).

### Simulated biparental population

In the simulated biparental populations, cross-validated predictive abilities in the initial doubled haploid population were high in all scenarios (0.69, 0.68, 0.52 and 0.61; Table 1), as expected when every mutation is genotyped. GS had the clearest advantage over MAS when the mutation rate was high and favorable and unfavorable effects were equally frequent (Figure 4). MAS is ineffective in that situation because it can select only one mutation. Details of each scenario are described below.

**Figure 4.**
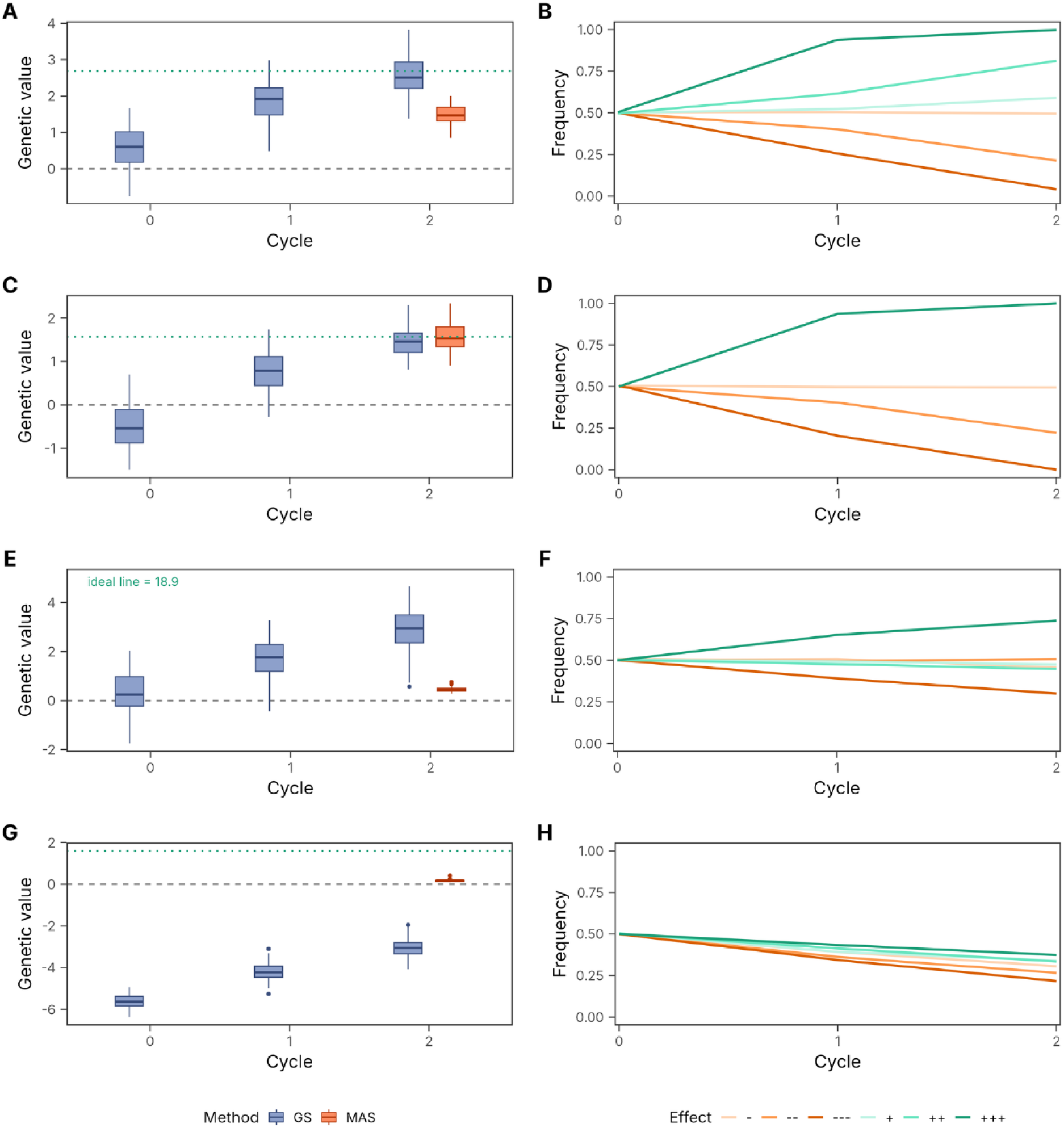
Genomic selection in simulated bi-parental populations (mutant x WT). Genetic values (A, C, E, G) of doubled haploid populations from a cross between an advanced mutant and its wild type (WT) over two cycles of selection (phenotypic selection in cycle 0, GS in cycle 1), and after MAS for the largest favorable mutation; boxes summarise 100 replicates. The dashed line marks the WT (genetic value 0). The dotted green line marks the ideal line carrying all favorable mutations; where it falls outside the panel its value is given as a label. Mutant allele frequencies (B, D, F, H) are shown for representative mutations at fixed effect ranks (the largest, a middle and the smallest unfavorable (−) and favorable (+) effects), averaged over replicates. [A–B] Low mutation rate (10 mutations) and equal proportions of favorable and unfavorable effects. [C–D] Low mutation rate and higher proportion of unfavorable effects. [E–F] High mutation rate (1,000 mutations) and equal proportions. [G–H] High mutation rate and higher proportion of unfavorable effects.

**Table 1.** Predictive ability and genetic values in simulated biparental populations (mutant. × WT).

| Mutation rate | Proportion favorable | Predictive ability, mean (SD) | Range | GS, cycle 2 | MAS | Ideal line |
| --- | --- | --- | --- | --- | --- | --- |
| Low (10 mutations) | 0.5 | 0.69 (0.03) | 0.61–0.76 | 2.53 | 1.50 | 2.68 |
| Low (10 mutations) | 0.1 | 0.68 (0.04) | 0.61–0.82 | 1.45 | 1.57 | 1.57 |
| High (1,000 mutations) | 0.5 | 0.52 (0.06) | 0.32–0.65 | 2.84 | 0.45 | 18.95 |
| High (1,000 mutations) | 0.1 | 0.61 (0.05) | 0.50–0.72 | –3.06 | 0.17 | 1.60 |

Predictive ability is the correlation between gEBV and phenotype in five-fold cross-validation of the initial 200 doubled haploid lines (mean, SD and range over 100 replicates). Genetic values are means over 100 replicates. The WT has a genetic value of 0, and the ideal line carries all favorable mutations and none of the unfavorable ones.

Both GS and MAS produce improvement in the simulation scenario with low mutation rate and equal proportions of favorable and unfavorable mutation effects, but the improvement from GS is higher than that from MAS (2.53 vs 1.50 after two cycles; Figure 4A). The additional improvement with GS can be seen with the increase in the frequencies of alleles with small and moderate favorable effects that are ignored in MAS (Figure 4B). Even though MAS is more effective at eliminating alleles with small and moderate unfavorable effects through background selection, it cannot select multiple favorable alleles like GS. As the selection cycle goes on, most of the favorable alleles will reach fixation and most of the unfavorable alleles will be lost.

With a low mutation rate and mostly unfavorable effects, MAS gave a slightly higher genetic value than GS after two cycles (1.57 vs 1.45, and GS was lower in all 100 replicates; Figure 4C). There are 10 mutations and only one is favorable, so GS has no small or moderate favorable effects to exploit and converges on the MAS outcome, which in this scenario equals the ideal line (Figure 4D). Some small and moderate unfavorable effects also take more cycles to purge with GS than with background selection in MAS.

With a high mutation rate and equal proportions of favorable and unfavorable effects, GS gave a much greater improvement than MAS (2.84 vs 0.45 after two cycles; Figure 4E). The gain came mainly from raising the frequency of the largest favorable mutations and purging the largest unfavorable ones, while mutations of small effect stayed close to their starting frequency (Figure 4F). Much of the variation remained untapped: the ideal line carrying all favorable mutations would have a genetic value of 18.95, so GS captured 15% of this value after two cycles and 23% after five (4.35).

GS fell short of MAS only when the mutation rate was high and most effects were unfavorable (Figure 4G). The few favorable mutations are tightly linked to many unfavorable ones, so GS could not raise their frequencies (Figure 4H), and the population mean stayed below the WT even after five cycles (−1.70, against 0 for the WT). MAS, which keeps a single favorable mutation and removes all others, was only marginally better than the WT (0.17). A mutant line with a heavy load of unfavorable mutations therefore offers little to either strategy.

### Simulated multiparental population

In the simulated multiparental populations, predictive abilities at cycle 0 were higher in populations segregating for both natural variants and mutations (Pop 2) than in those segregating for natural variants only (Pop 1) in every scenario (Figure 5A), partly because Pop 2 has the larger total genetic variance. Whether mutations added genetic gain depended on the mutational variance (Figure 5B–D, S3). When V_mut was large, Pop 2 gained more than Pop 1 in all four scenarios. When V_mut was small and V_nat large, the advantage was small (gain over five cycles 5.10 vs 4.85 with PF = 0.5, and 4.80 vs 4.62 with PF = 0.1). Three key scenarios are described below.

**Figure 5.**
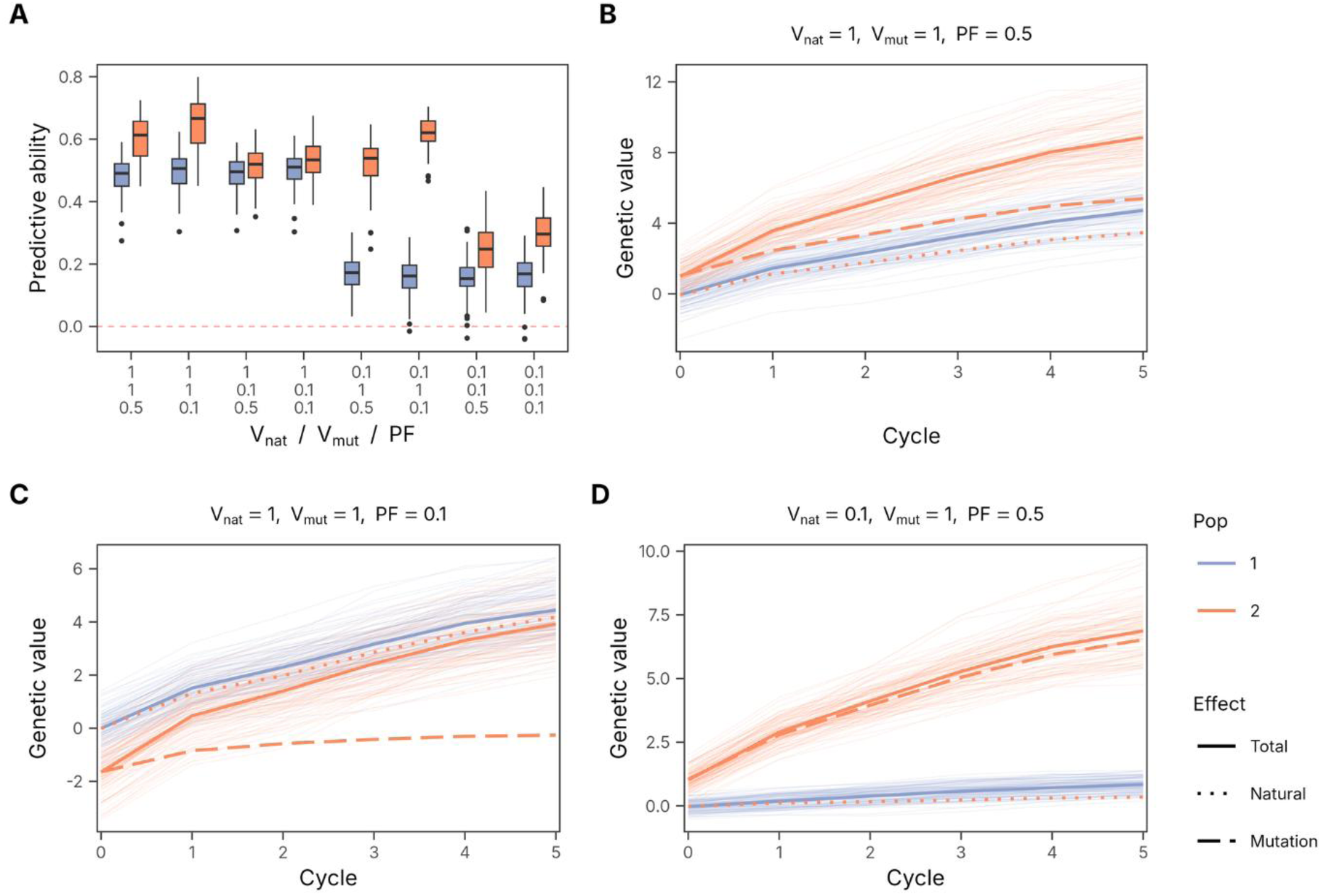
Genomic selection in simulated multiparental populations. [**A**] Predictive ability (correlation between gEBV and phenotype in five-fold cross-validation at cycle 0). [**B**–**D**] Mean genetic value over five cycles of selection (phenotypic selection in cycle 0, GS from cycle 1). Thin lines show the 100 replicates and thick lines their mean. Eight combinations of parameters were simulated: high (1.0) and low (0.1) natural genetic variance (V_nat_), high (1.0) and low (0.1) mutation genetic variance (V_mut_), and equal (0.5) and unequal (0.1) proportions of favorable mutation effects (PF). Pop 1 segregates for natural variants only; Pop 2 segregates for natural variants and mutations, and its genetic value is partitioned into natural and mutation components. [**B**] V_nat_ = 1.0, V*mut* = 1.0, PF = 0.5. [**C**] V_nat_ = 1.0, V_mut_ = 1.0, PF = 0.1. [**D**] V_nat_ = 0.1, V_mut_ = 1.0, PF = 0.5. All eight scenarios are shown in Figure S3.

With high natural and mutation genetic variance and equal proportions of favorable and unfavorable mutation effects, Pop 2 gained 7.87 over five cycles, against 4.77 for Pop 1 (Figure 5B). Its gain from natural variants (3.51) was smaller than that of Pop 1, but its additional gain from mutations (4.37) more than compensated, so its genetic value at cycle 5 was 1.9 times that of Pop 1.

With high natural and mutation genetic variance and mostly unfavorable mutation effects, Pop 2 started from a lower genetic value because its mutant founders carry a net unfavorable mutation load (−1.65 at cycle 0; Figure 5C). Its gain over five cycles was larger than that of Pop 1 (5.58 vs 4.46), but this did not fully make up the initial deficit by cycle 5 (3.92 vs 4.45).

With low natural genetic variance, high mutation genetic variance and equal proportions of favorable and unfavorable effects, Pop 2 gained 5.85 over five cycles, against 0.86 for Pop 1, and 94% of its gain came from mutations (Figure 5D). This scenario shows the clearest advantage of combining mutation breeding with GS in populations with limited genetic diversity.

## DISCUSSION

Applying GS directly across the independent lines of an advanced mutant panel does not test what it appears to test. Induced mutations are almost all private to one line, so their effects cannot be estimated from the other lines of a training set, and any predictive ability must come from variants shared among lines. This is what we observed. In wheat, 34 of the 190 lines were off-types, and within the true-to-type lines 97% of non-WT SNP calls were shared and formed chromosome segments in almost complete linkage disequilibrium, consistent with residual heterogeneity in the parental cultivar. In tomato, 97% of variant sites were private. In both species, shared variants alone reproduced the predictive abilities obtained with all markers, and private variants alone predicted nothing. Collapsing variants into genes or GO categories does not change this, because independent mutations rarely hit the same genes. Advanced mutant panels are therefore not suitable training populations for predicting induced variation. Apparent predictive ability in such panels should be checked against the WT and partitioned into shared and private variants before it is interpreted. The value of GS for mutation breeding lies instead in populations in which mutations are shared among relatives, which crossing provides.

The simulated biparental populations show that polygenic induced variation becomes accessible to GS once mutations segregate. When mutations were numerous and favorable and unfavorable effects were balanced, GS accumulated favorable mutations across cycles and outperformed MAS for the largest mutation by a factor of six after two cycles. Conventional MAS fixes a single mutation identified by mapping approaches such as MutMap (Abe et al 2012) or MutMap+ (Fekih et al 2013) and removes the rest by background selection, so it discards most of the variation created by mutagenesis. Two limits are also clear. First, linkage among many mutations restricts progress: GS captured only 15–23% of the value of the ideal line within five cycles, so larger populations, more recombination or haplotype stacking could release more. Second, when unfavorable mutations dominate, neither GS nor MAS improves meaningfully on the WT, so mutant lines with heavy loads should be screened out before crossing. These results assume that every mutation is genotyped and that no non-causal markers are present, which is a best case. In practice, marker density and mutation detection will reduce accuracy.

The simulated multiparental populations show how mutation breeding could feed pre-breeding. Crossing advanced mutants derived from different founders lets linkage disequilibrium between natural variants and mutations carry mutation effects into GS models without sequencing each mutant. When mutation genetic variance was large and favorable and unfavorable effects were balanced, populations with mutant founders gained 1.7 times as much over five cycles as populations without mutations at high natural variance, and 6.8 times as much at low natural variance, where mutations contributed 94% of the gain. The latter is the situation of breeding programmes with a narrow genetic base. When mutation variance was small relative to natural variance, mutations added little (4–5% more gain with V_nat = 1), because selection on a small mutational component competes with selection on natural variants. Whether mutagenesis is worthwhile therefore depends on the ratio of mutational to standing variance, which is rarely measured for induced mutations and should be estimated in mutant-derived populations. Polygenic mutations are pervasive, and these results support a shift in mutation breeding from searching for single mutations toward continuous germplasm enrichment with pools of favorable mutations, provided that mutational variance is substantial.

Moving forward, emerging technologies can be used to enhance the integration of mutation breeding with genomic selection. First, phenomic selection with high-throughput phenotyping (Rincent et al 2018) reduces phenotyping cost and increases training population size in early mutant generations. Second, long-read sequencing technologies enables accurate profiling of complex structural variants, copy number variants and large indels induced by mutagens and improves the resolution of genome-wide mutations (McCord et al 2022). Third, multiplex gene editing tools allow direct induction of polygenic variation across targeted pathways while bypassing unfavorable mutations (Chen et al 2019). While strategies for selecting against evolutionary deleterious mutations have been explored (Ramstein and Buckler 2022; Long et al 2023), the application of GS in pre-breeding pipelines offer a proactive framework for broadening genetic diversity in breeding populations to better meet the challenges of climate resilience and global food security (Xiong et al 2022; Cooper and Messina 2023).

## DECLARATIONS

### Data availability

The wheat genotype and phenotype data are from Guo et al (2019) and the tomato data from Shikata et al (2016) and Yano et al (2019) (TOMATOMA, https://tomatoma.nbrp.jp). The processed input files used here are provided with the code.

### Code availability

All R scripts used for the analyses and simulations are available at Zenodo https://zenodo.org/records/23689223.

### Funding

IAEA Coordinated Research Project D24015 research contract; Scottish Government RESAS & SRUC funding.

### Competing interests

IM was a member of the Editorial Board of Theoretical and Applied Genetics. The other authors declare no competing interests.

### Author contributions

CJY, IM, RS and WP conceived the study. IM led the project and secured funding through the IAEA Coordinated Research Project. CJY performed the analyses and simulations and wrote the first draft. RS reviewed the code, re-ran the analyses and revised the manuscript. WP supervised the project and revised the manuscript. CJY, RS and WP read and approved the final manuscript. IM died before the manuscript was completed; his family approved his inclusion as an author.

## Notes

### Competing Interest Statement

The authors have declared no competing interest.

