## Supplementary for "Mutation breeding for quantitative traits in the era of genomic selection"

**Supplementary File**

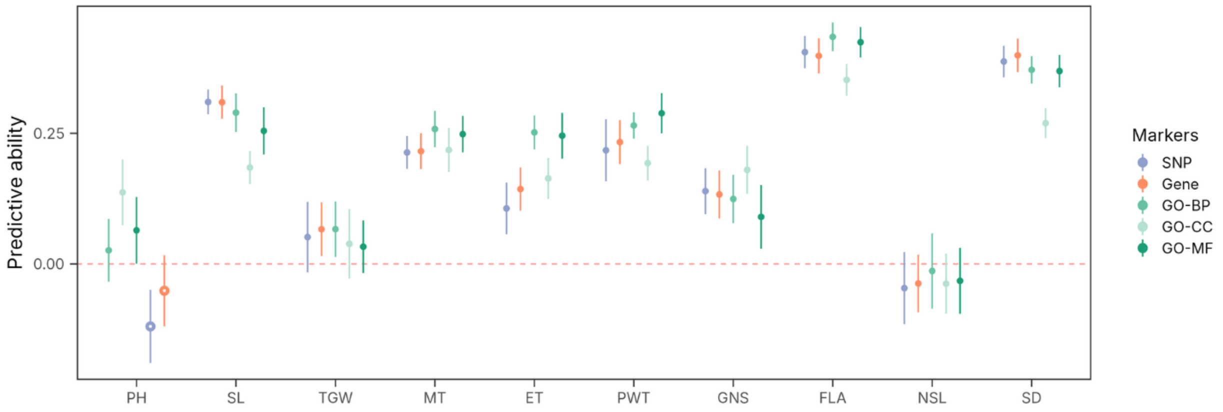


**Figure S1. Prediction with gene and Gene Ontology markers in the advanced wheat mutant population**

Predictive ability (mean ± SD over 20 repeats of five-fold cross-validation) in the 156 true-to-type wheat lines, with genomic relationship matrices built from SNPs, from SNPs collapsed into genes, and from genes collapsed into Gene Ontology biological process (GO-BP), cellular component (GO-CC) and molecular function (GO-MF) terms. Open circles mark estimates for which the genetic variance was estimated at zero in more than half of the 100 folds.

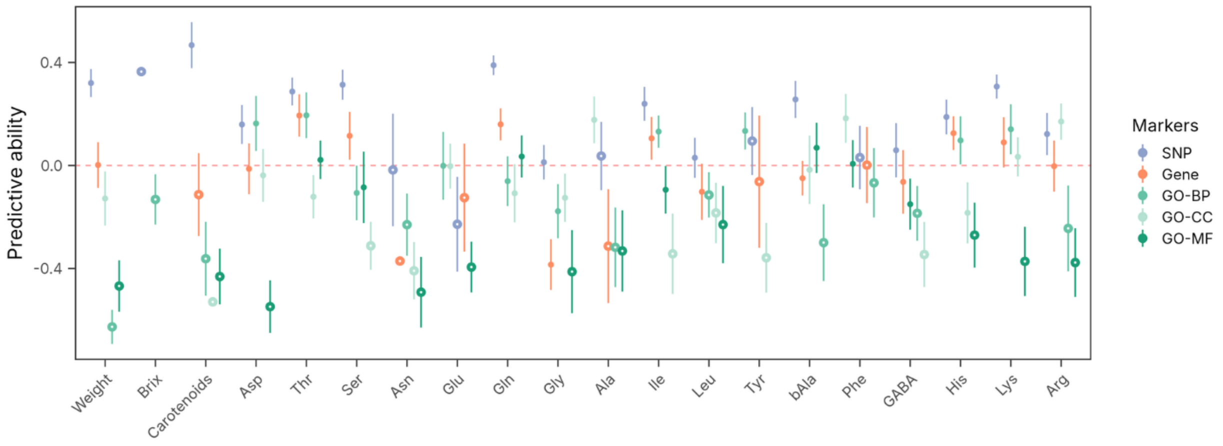


**Figure S2. Prediction accuracies in advanced tomato mutant population**

Predictive ability (mean ± SD over 20 repeats of five-fold cross-validation) for all 20 traits in the 74 tomato lines, with genomic relationship matrices built from variant sites, from sites collapsed into genes, and from genes collapsed into GO-BP, GO-CC and GO-MF terms. Open circles as in Figure S1.

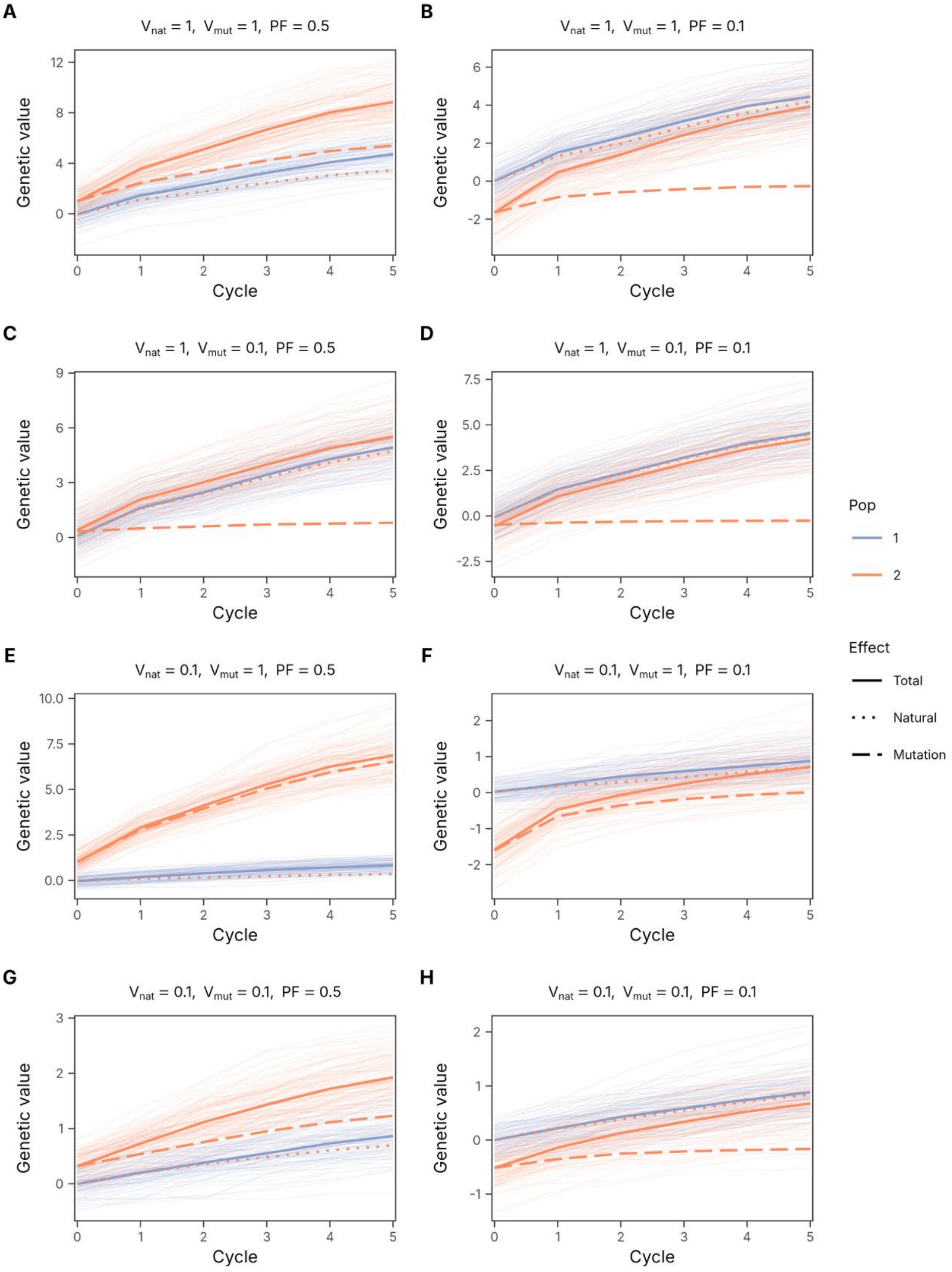


**Figure S3. Genetic gains in simulated multiparental populations.**

Thin lines show the 100 replicates and thick lines their mean. There are 8 different combinations of simulation parameters, including high (1.0) and low (0.1) natural genetic variance ($V_{nat}$), high (1.0) and low (0.1) mutation genetic variance ($V_{mut}$), and equal (0.5) and unequal (0.1) proportions of favorable and unfavorable mutation effects (PF). For each combination, there are two simulated multiparental populations where Pop 1 segregates for only natural genetic variants and Pop 2 segregates for natural genetic variants and mutations. Specifically for Pop 2, the total genetic gains are further partitioned into natural and mutation sources. [**A**] $V_{nat}$=1.0, $V_{mut}$=1.0, PF=0.5, [**B**] $V_{nat}$=1.0, $V_{mut}$=1.0, PF=0.1, [**C**] $V_{nat}$=1.0, $V_{mut}$=0.1, PF=0.5, [**D**] $V_{nat}$=1.0, $V_{mut}$=0.1, PF=0.1, [**E**] $V_{nat}$=0.1, $V_{mut}$=1.0, PF=0.5, [**F**] $V_{nat}$=0.1, $V_{mut}$=1.0, PF=0.1, [**G**] $V_{nat}$=0.1, $V_{mut}$=0.1, PF=0.5, [**H**] $V_{nat}$=0.1, $V_{mut}$=0.1, PF=0.1.
